# ADAGE analysis of publicly available gene expression data collections illuminates *Pseudomonas aeruginosa-host* interactions

**DOI:** 10.1101/030650

**Authors:** Jie Tan, John H. Hammond, Deborah A. Hogan, Casey S. Greene

## Abstract

The growth in genome-scale assays of gene expression for different species in publicly available databases presents new opportunities for computational methods that aid in hypothesis generation and biological interpretation of these data. Here, we present an unsupervised machine-learning approach, ADAGE (<u>A</u>nalysis using <u>D</u>enoising <u>A</u>utoencoders of <u>G</u>ene <u>E</u>xpression) and apply it to the interpretation of all of the publicly available gene expression data for *Pseudomonas aeruginosa,* an important opportunistic bacterial pathogen. In post-hoc positive control analyses using curated knowledge, the *P. aeruginosa* ADAGE model found that co-operonic genes often participated in similar processes and accurately predicted which genes had similar functions. By analyzing newly generated data and previously published microarray and RNA-seq data, the ADAGE model identified gene expression differences between strains, modeled the cellular response to low oxygen, and predicted the involvement of biological processes despite low level expression differences in directly involved genes. Comparison of ADAGE with PCA and ICA revealed that ADAGE extracts distinct signals. We provide the ADAGE model with analysis of all publicly available *P. aeruginosa* GeneChip experiments, and we provide open source code for use in other species and settings.

## Importance

There has been a rapid increase in the availability of genome-scale data sets that examine RNA expression in diverse bacterial and eukaryotic species. Thus, we can greatly benefit from analytical methods that do not rely on existing biological knowledge for model construction. Our ADAGE method integrates such data without requiring gene class or experiment labeling, making its application to any large gene expression compendium practical. The *Pseudomonas aeruginosa* ADAGE model was derived from a diverse set of publicly available experiments without any prespecified biological knowledge, and this model was accurate and predictive. The ADAGE results that we provide for the complete *P. aeruginosa* GeneChip compendium can be used by researchers studying *P. aeruginosa* and the provided source code will allow ADAGE to be applied to other species.

## Introduction

Modern biomedical research routinely generates rich datasets measuring genome-wide gene expression, and advances in sequencing technology have dramatically reduced the cost and increased the use of genome-wide assays of gene expression (1–7). Many methods exist to identify important signals from data generated within a single experiment, e.g. clustering (8–10) or differential expression analysis (11), but integrative analyses across many datasets are more challenging, particularly in microbial systems in which many different conditions are assessed. In well-studied species, integrative analyses of gene expression often employ supervised methods that leverage prior knowledge to extract information from noisy data present in large publicly available datasets (12–14). In less well-studied organisms, the task of large-scale gene expression analysis is more challenging (15, 16) due to limited information about gene function and the absence of prior knowledge about the organism’s biology. As the accumulation of data exceeds curation, particularly in non-model organisms, new unbiased approaches to reveal biological patterns are required.

Deep learning algorithms have transformed how underlying explanatory factors are extracted from diverse large-scale unlabeled datasets (17). Denoising Autoencoders (DAs) (18), examples of one form of deep learning, extract important signals and construct representative features, referred to as nodes, by training models to remove noise that is intentionally added to input data. DAs successfully recognize hand-written digits (18), spoken words (19), and the sentiment of Amazon reviews (20). Because the DA’s learning objective is defined entirely by the data, this algorithm can extract meaningful features without requiring prior knowledge, which makes DAs well suited to the challenge of data integration for non-model organisms.

Here, we report the development of an approach termed <u>A</u>nalysis using <u>D</u>enoising <u>A</u>utoencoders of <u>G</u>ene <u>E</u>xpression (ADAGE) capable of integrating diverse gene expression data to aid in the interpretation of existing and new experiments. Using an unsupervised machine learning approach, the community-wide *Pseudomonas aeruginosa* gene expression data were integrated to create an ADAGE model that captures patterns that correspond to biological states or processes in gene expression data. In our analysis, each dataset is interpreted in terms of the activity of fifty distinct nodes, with each node being influenced by different sets of genes. In positive control analyses, we found that co-operonic genes were preferentially linked to common nodes, and that genes with similar KEGG functions had similar gene-node relationships across the model. More interestingly, ADAGE extracts certain nodes representing recognizable identities with predictive value. Additionally, we show that ADAGE is capable of revealing subtle but biologically meaningful signals within existing datasets. We compared ADAGE with existing popular feature construction approaches including principal component analysis (PCA) and independent component analysis (ICA). The features captured by ADAGE were not fully extracted by either PCA or ICA. The unsupervised ADAGE approach can be applied to any large publicly available gene expression compendium or newly generated gene expression data to characterize genomic and transcriptional features.

## Results

### Construction of an ADAGE Model for *P. aeruginosa*

To build the ADAGE model for the analysis of *P. aeruginosa* gene expression, we focused on expression profiling performed using Affymetrix GeneChips because of the uniform gene nomenclature. All *P. aeruginosa* GeneChip expression data were downloaded from the ArrayExpress database (21), and this resulted in a compendium of 950 arrays from 109 experiments (Supplemental File 1). We constructed an ADAGE model from the compendium by first adding random noise to the input data and then training a neural network with hidden nodes that were able to remove added noise to reconstruct the initial data (Figure 1, details in Methods). The process of adding noise improves the robustness of constructed features and consequently the resulting models (22, 23). The resultant network was designed to contain fifty nodes, and within each node all *P. aeruginosa* strain PAO1 genes were assigned weights that reflected the contribution of each gene to the activity of each node (weight vectors provided in Supplemental File 2). A model with 50 nodes was chosen to balance reconstruction error with the need to manually interpret the ADAGE model, and our subsequent analyses demonstrate that networks of this size are capable of adequately extracting major global transcriptional patterns (Figure 1).

**Figure 1:**
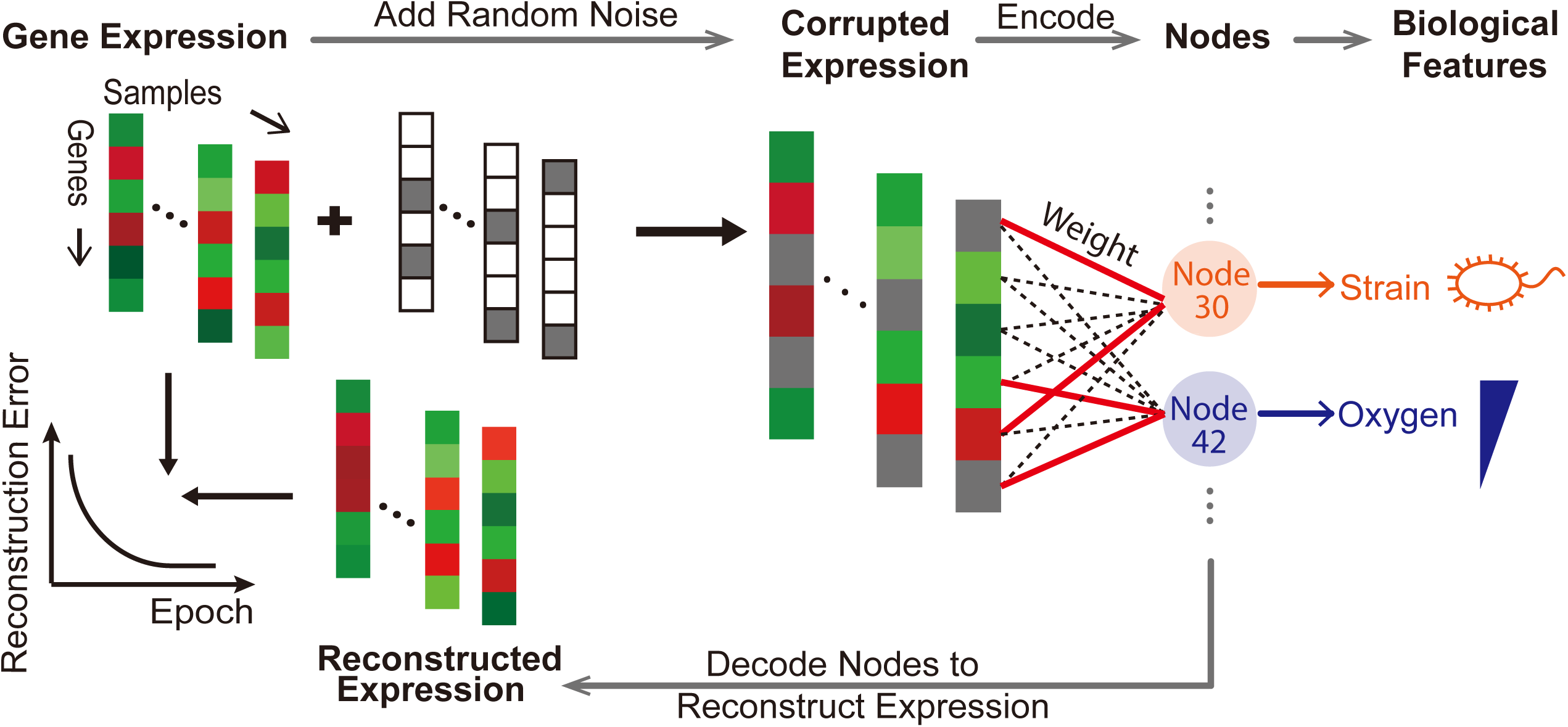
Analysis using Denoising Autoencoders of Gene Expression (ADAGE). For one sample in the expression compendium (one column in the figure with red or green colors representing expression values of various genes), random noise is first added to the expression value. The corrupted expression values are then encoded into 50 nodes through a gene-to-node weight matrix, which connects each gene to each node. A red solid line represents a high-weight relationship between a gene and a node, indicating that gene has a stronger influence on the node’s activity than other genes (connected by black dotted lines). Node activities derived from this sample are decoded back into reconstructed expression values through the same weight matrix. Samples in the compendium are trained through the encoding and decoding steps with the goal of minimizing differences between initial expression values and reconstructed expression values. The resulting ADAGE model constructs nodes from genomic measurements that can be interpreted as biologically meaningful features such as genome divergence among strains and transcriptional responses to oxygen abundance.

All genes were connected to each node by a weight vector, and the contributions, or gene weights, within a node were distributed symmetrically and approximately centered at 0. These weights approximately resembled a normal distribution in which a small proportion of genes provided high positive or high negative weights to that node (Figure 2A). We refer to genes that were outside of two standard deviations as high-weight (HW) genes for that node (red regions of Figure 2A). In the ADAGE model, 4029 genes (72.6% of the *P. aeruginosa* genome) are HW in at least one node (Figure 2B, outermost ring and Supplemental File 3), and 229 (4.1%) genes are HW in ten or more nodes. It is important to convey that nodes differed in terms of the identities of the HW genes. The innermost rings of Figure 2B show the HW genes in two example nodes (node 42 and 30 respectively from outside to inside), which we identified as representative of anaerobic response and strain specificity as discussed in more detail below.

**Figure 2:**
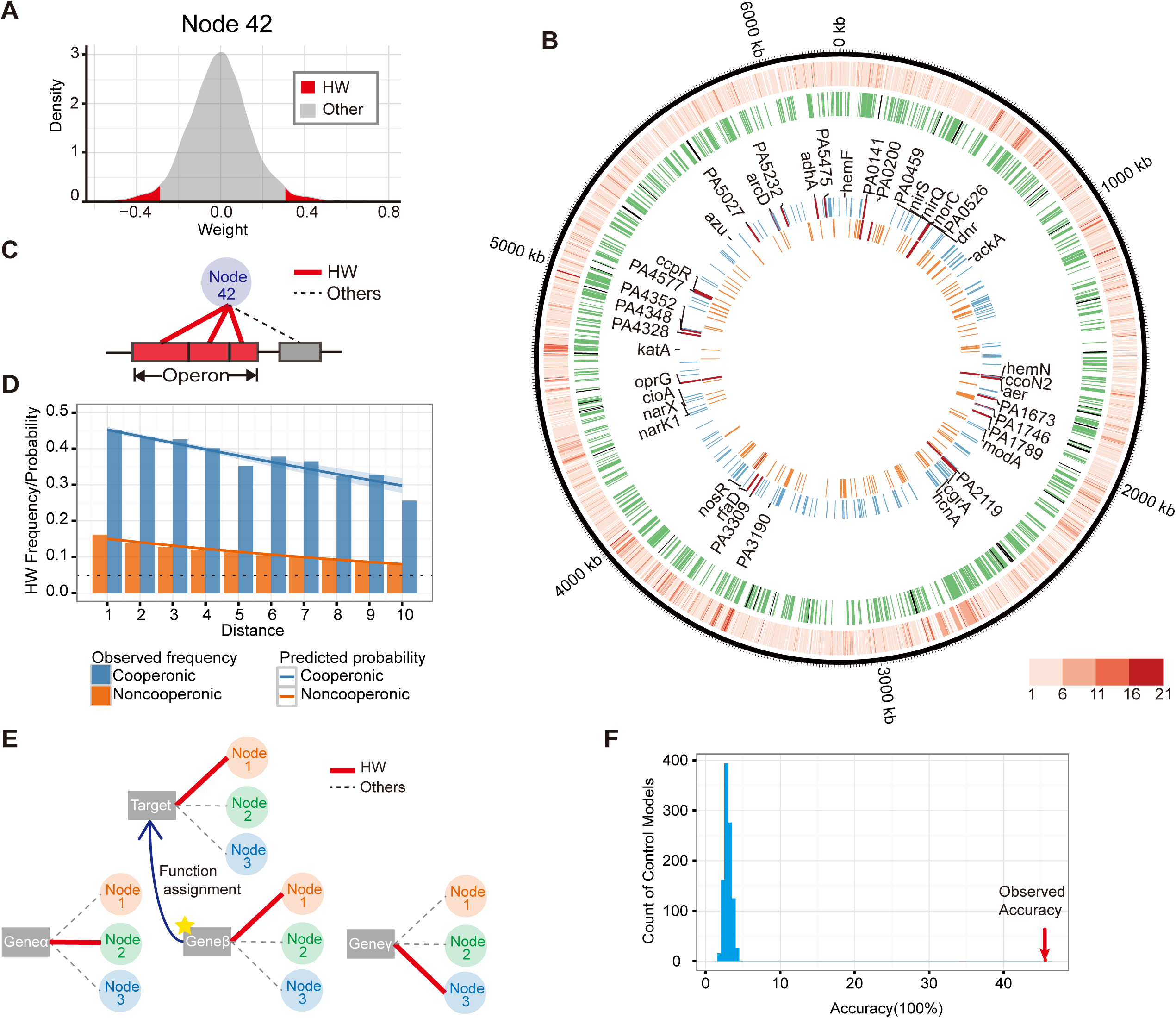
ADAGE weights reflected gene’s common regulatory and process features. (**A**) HW genes defined in ADAGE. The distribution of edge weights that connect genes to each node, e.g. Node 42 shown here, is approximately normal. HW genes in a node were defined as genes whose weights were more than two standard deviations from the mean (shown in red). (**B**) The contributions of individual genes and operons to the ADAGE model. The outermost ring shows those genes that are HW within at least one node. The color intensity reflects how many nodes (ranging from 1 to 21) a gene was connected to at a high weight. The second to outermost circle shows operons that were significantly associated with at least one node (green bars) and those that were not significantly associated with any of the 50 nodes (black bars). The inner two circles represent the HW genes in nodes 42 and 30 (outside to inside). The labeled genes are those identified in Trunk et al. (37) and Jackson et al. (36) as regulated by Anr and they are colored in red among HW genes of nodes 30 or 42. The HW genes in Node 42 overlapped extensively with Anr regulated genes, which suggested that Node 42 captured the regulatory signature of Anr. (**C**, **D**) ADAGE captured principles of bacterial genome organization. (**C**) In the bacterial genome, genes are arranged into operons, which the ADAGE model recognized by connecting co-operonic genes to a shared node. (**D**) The regulatory role of genome positioning in *P. aeruginosa* was captured by ADAGE. A logistic regression analysis revealed that co-operonic genes (blue line shows model; bars show observed) were more likely to be co-HW genes than non-co-operonic genes (red). As the number of genes between two genes on the chromosome increased, they were less likely to be co-HW. The black dotted line represents the background frequency of HW genes across all nodes. (**E, F**) ADAGE captured gene’s functional features. (**E**) We found the closest neighbor of a target gene based on the Euclidean distance between the weight vectors connecting each gene to 50 nodes and assigned the closest neighbor gene’s function to the target gene. (**F**) The accuracy of gene function assignment using ADAGE model (45%, pointed by the red arrow) was much higher than the accuracies achieved with 1000 randomly permuted control models (distribution shown in blue). Here we considered a function assignment as positive if one or more functions assigned by the closest neighbor gene match the target gene’s annotations.

### Operonic co-membership and spatial proximity reflect gene-node relationships

Bacterial operons, by definition, contain genes that are co-expressed, though genes within an operon may be transcribed by multiple promoters. To determine if genes within operons shared similar node relationships, Gene Set Enrichment Analysis of ADAGE weights using operon annotations from DOOR (24) were performed and results showed that, in total, 92.9% of co-operonic genes were significantly associated with at least one common node (Figure 2B, second to outermost circle; operons significantly associated with at least one node are colored in green and unassociated operons are in black). The operons significantly associated with each node are shown in Supplemental File 4. As an extension of the above analysis of operon-node relationships, we also predicted co-operonic genes would be HW to the same nodes (Figure 2C). To test this prediction, we fit a logistic regression model that predicted whether a gene was likely to be HW in a given node based on whether co-operonic genes were also HW to that node. As predicted, genes co-operonic with a HW gene had a 4.6 times higher odds of being HW in the same node. In addition, we determined if genes were more likely to be HW to the same node if they were spatially proximal (e.g. a small number of intervening genes) even if they were not co-operonic. Again, as expected, the odds of two adjacent genes being HW to the same node were higher for co-operonic genes than for non-co-operonic genes (Figure 2D). Interestingly, for both co-operonic and non-co-operonic genes, every additional intervening gene between two genes decreased the odds of them being HW to the same node by a factor of 0.9 (Figure 2D) indicating links between proximity and co-expression. This trend could reflect that genes within a pathway are often physically close or that there are other regional factors that affect local gene expression in a coordinated way.

### Genes within a common KEGG pathway share node relationships

To further test the biological relevance of the ADAGE model, we predicted that genes involved in the same pathway would have similar gene-node relationships. We tested this prediction using post hoc analysis of the ADAGE model via the Kyoto Encyclopedia of Genes and Genomes (KEGG) (25). To do so, we employed a straightforward algorithm in which a target gene is assigned the KEGG function of its closest “neighbor” based on ADAGE weights; the neighbors for each gene were determined by calculating the Euclidean distance between each gene’s connections to all nodes for every gene pair (Figure 2E). If a gene’s predicted KEGG functions based on the functions of its closest ADAGE model neighbor matched at least one of its actual KEGG annotations, it was considered a positive. If no KEGG annotations matched, it was considered a negative. We used this approach because our goal is to evaluate the model itself. Though more complex techniques would be likely to provide superior predictions of function, they would not be as useful for direct evaluation of the underlying model. We observed a high accuracy of gene-function assignment (45%) with the ADAGE model. As a control, the identical algorithm was applied to 1000 control models, where we randomly permuted gene identifiers. In these tests, the mean accuracy was 3% with no permuted model achieving greater than 5% accuracy (Figure 2F). We also evaluated a more stringent definition of correct assignment that requires all predicted and annotated functions to match and observed consistent results. In this analysis, 34% of gene-function assignments were correct when the ADAGE model was used while less than 3% of assignments were correct when randomly generated models were used. These analyses suggest that ADAGE grouping of genes into nodes based on expression across the *Pseudomonas* Gene Chip compendium identified biologically-relevant relationships between genes.

### ADAGE Recognizes Genomic Differences between Strains

We predicted that the HW genes within a node were likely grouped together because they were related in their expression patterns across many datasets in the compendium. Visual inspection of the lists of HW genes (Supplemental File 3) revealed that several nodes contained genes that are known to vary between strains (LPS, flagellin, pili, etc.) (26). Because the *P. aeruginosa* compendium includes experiments performed on different *P. aeruginosa* strains, we sought to determine if strain-specific signatures were represented in the ADAGE model. To do this, we isolated DNA from two well-studied strains of *P. aeruginosa,* PAO1 (27) and PA14 (28) and performed a DNA hybridization experiment. Hybridization to the *Pseudomonas* GeneChip yielded a profile in which a small number of ADAGE nodes were highly differentially active (Figure 3A). In the three most differentially active nodes (30, 33 and 25 in order of the difference magnitude), we found many genes that are known to associate with strain-to-strain differences within the species. We focused our further analyses on Node 30, which exhibited the largest difference in activity between PA14 and PAO1. A functional enrichment analysis by Gene Ontology (GO) (29) and KEGG (25) terms found that the HW genes in Node 30 included those associated with the surface exposed portions of type IV pili *(pilA, pilC, pilV, pilW, pilY1, pilY2)* and the flagellum *(flgK, flgL, fliC, fliD)* (Supplemental Table 1). Importantly, the ADAGE model precisely identified specific genes within pili and flagellar gene clusters as contributing strongly to the identity of Node 30. For example, two genes involved in pili biosynthesis, *pilA* and *pilC,* were among the HW genes in Node 30 while the adjacent genes *pilB* and *pilD* were not (Figure 3C for gene relationships). We performed an alignment and pairwise comparison of the *pilABCD* coding sequences from thirteen sequenced *P. aeruginosa* strains. This analysis revealed that *pilA* and *pilC* were strikingly more divergent than either *pilB* and *pilD,* or than other adjacent genes *nadC* and *coaE* (Figure 3C and 3D). In fact, *pilA,* had the highest weight in Node 30 and was the most divergent gene of those analyzed (Figure 3C and 3D). A similar trend was observed for the flagellum-associated genes; the five HW flagellar genes in Node 30 varied in sequence across strains (Figure 3C), but two adjacent genes, *fleQ* and *fleS* were highly conserved and were not HW in Node 30 (Figure 3C). In further support of the hypothesis that the activity of Node 30 identifies strain differences, this node contains other strain-specific genes, including those involved in LPS biosynthesis *(wbpA, wbpB, wbpD, wbpE, wbpG, wbpH, wbpI, wbpJ, wbpK, wbpL,* and *wzz),* a putative type I restriction/modification system (PA2730-PA2736), and pyoverdine biosynthesis. The genes encoding bacteriophage Pf4 (PA0717-PA0734), which is only found in certain strains of *P. aeruginosa* (30), as well as the highly strain-specific R-pyocins (PA0621-PA0648) (31), were also among the HW genes in Node 30. HW genes in the strain-differentiating nodes also included genes that were either unique to PAO1 (PA3501-3504) or only found in a subset of the *P. aeruginosa* strains with published genomes, such as PA0202-6, which encode putative transporter genes.

**Figure 3:**
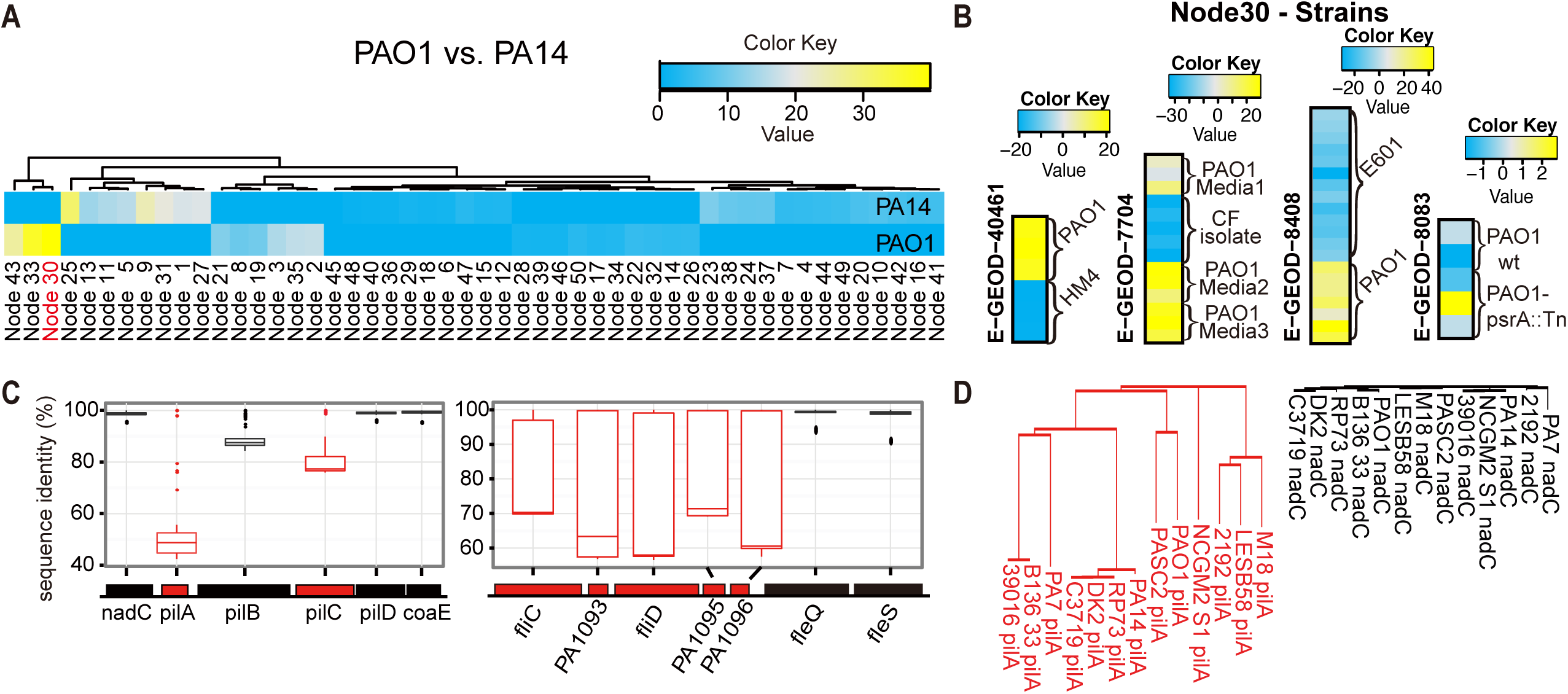
ADAGE extracted features that represent sequence differences between strains. (**A**) ADAGE node activity heatmap analysis of the genome hybridization of *P. aeruginosa* strains PA14 and PAO1 to Affymetrix *Pseudomonas* GeneChip. The heatmap shows the ADAGE activity differences of each node in each strain when the genome hybridization data were analyzed. The activity values of Node 30 most strongly differentiated strains PAO1 and PA14. (B) Analysis of mean centered Node 30 activity values clearly distinguished strain PAO1 from other *P. aeruginosa* strains (HM4, E601) or uncharacterized CF strains in RNA expression experiments. For each heatmap, the complete range of mean-centered node activities is used to generate the color range. The range of activity values in experiments that compare strains is at least 40 (-20 to 20) where as the range of Node 30 activities was much less in a dataset in which only PAO1 or its derivatives were analyzed. (**C**) Pair-wise percent sequence identities between orthologous genes from 13 *P. aeruginosa* strains. Two operons that contain the top 5 HW genes in Node 30 are analyzed. HW genes are colored in red. (**D**) Phylogenetic trees of *pilA* (most HW gene) and *nadC* (*pilA*’s immediate neighbor). The two trees share the same distance per branch length unit and the longer branch represents more genetic changes.

To determine if Node 30 activity differed in gene expression experiments that made comparisons across strains, we identified published experiments within the *P. aeruginosa* GeneChip compendium in which multiple strains were measured. This analysis found that Node 30 was indeed differentially active in experiments that included the comparison of different *P. aeruginosa* strains (E-GEOD-40461, E-GEOD-7704 (32), E-GEOD-8408 (33) in Figure 3B). In the dataset E-GEOD-7704, Node 30 clearly indicates the differences between lab strain PAO1 and clinical Cystic Fibrosis (CF) strains. In contrast, Node 30 activity did not differ in a control experiment using the same strain (E-GEOD-8083 (34) in Figure 3B, the small range in color key indicates no difference among samples from the same strain.). In the discussion, we address the need for a community-wide allele nomenclature for variable genes to support these types of analyses across datasets and strains when presence/absence is not the read out for strain variation.

### ADAGE Node Activities Reflect Transcriptional Responses

To further test whether the node-based ADAGE model identified biological states in gene expression data, we analyzed experiments in which *P. aeruginosa* wild-type cells were compared to cells that lacked the transcription factor Anr. Anr is active in low oxygen environments and regulates the cellular response to oxygen limitation (35) and other virulence related processes (36). We found that Node 42 was most significantly enriched for HW genes known to be regulated by Anr (FDR q value = 4.24e-31). The Anr regulated genes used in this analysis (36, 37) are listed in Supplemental Table 2. To investigate whether ADAGE could reliably extract an Anr signal from the compendium, we built 100 ADAGE models with different random seeds and repeated the enrichment test. Across the 100 models, 80 contained nodes as significant or more significant than Node 42 in the current model. All 100 models contained a node significantly associated with Anr targets. This indicates that ADAGE robustly identified strong transcriptional patterns across independent applications of the algorithm.

We examined the activity of Node 42 in two datasets that compared wild type and Δ*anr* strains (E-GEOD-17179 (37) and E-GEOD-17296 (38) in Figure 4A). Node 42 showed low activity (blue) in the *anr* mutant, even in low oxygen conditions that would otherwise be expected to activate Anr (grey). In addition, we also evaluated the activity of Node 42 in datasets where the responses to different oxygen concentrations were compared and found that again, the activity of Node 42 was modulated by oxygen availability (E-GEOD-33160 (39) and E-GEOD-52445 (40) in Figure 4A). E-GEOD-52445 is a high-time-resolution analysis of *P. aeruginosa* transiting from high oxygen tension to low oxygen tension and then reverse (40). Node 42 activity values gradually increased as oxygen levels decreased in the dataset E-GEOD-52445, and the restoration of oxygen was concomitant with a striking decrease in Node 42 activity. The activity pattern for Node 42 in all of the datasets in the compendium can be viewed at the url: http://adage.greenelab.com/Paeruginosa-da/node42/index.html.

**Figure 4:**
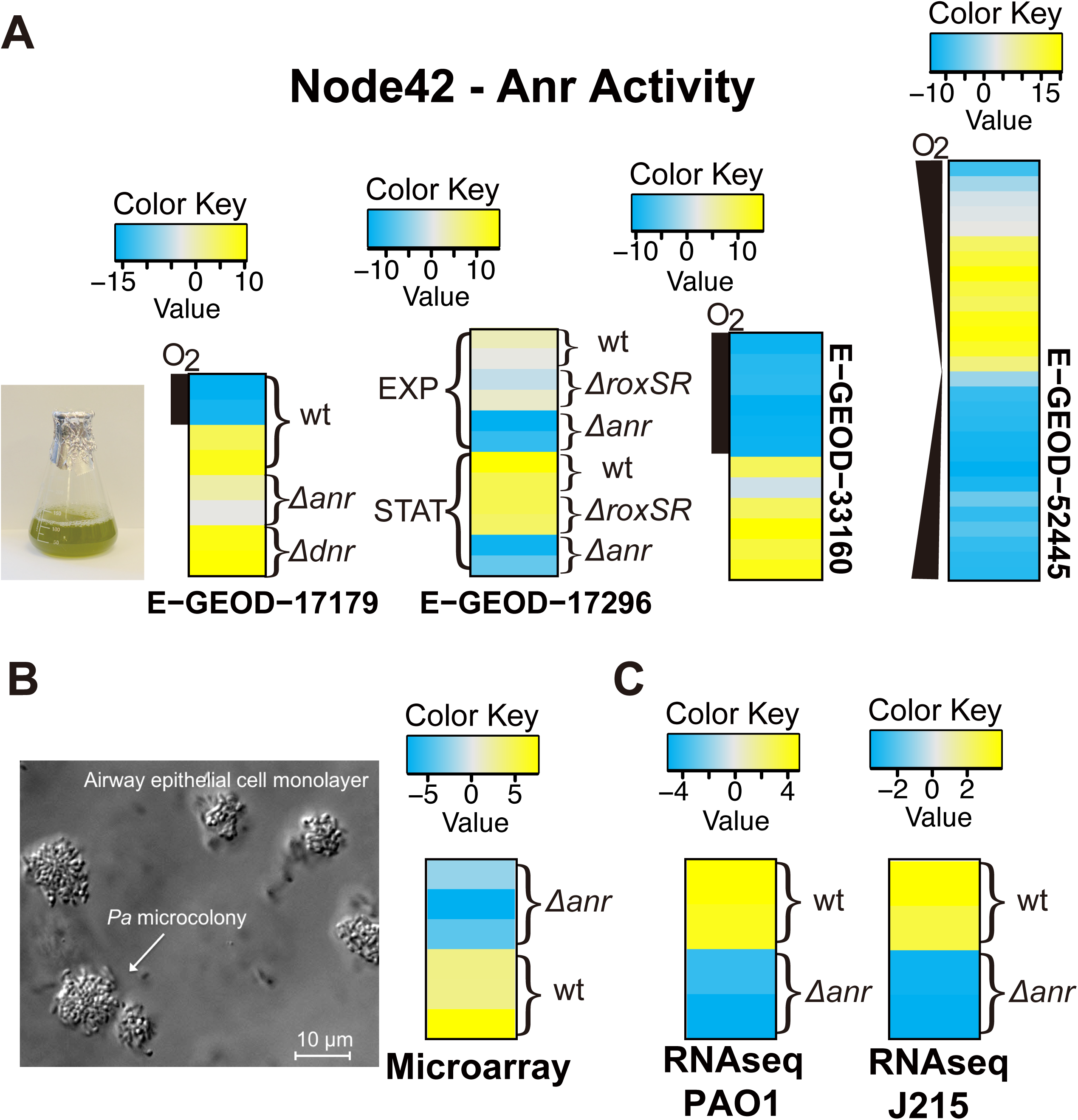
Node 42 reflected Anr activity in both existing and new experiments. (**A**) Mean-centered activity heatmaps of Node 42 for four datasets grown in liquid culture that examined *anr* mutants or altered oxygen levels. In E-GEOD-17179 and E-GEOD-17296, the low activity value (blue or grey in anaerobic condition) of Node 42 corresponded to *anr* deletion, and similar effects were not observed for *dnr* or *roxSR* mutants. In E-GEOD-17179 and E-GEOD-33160, the oxygen bar represents whether or not *P. aeruginosa* is in an aerobic or anaerobic environment. E-GEOD-52445 is a high-time-resolution experiment transiting from high oxygen tension to low oxygen tension, which is then reversed. The activity value of Node 42 was negatively correlated to the oxygen abundance in the microbes’ living environment. These results reflected Anr’s role as a transcriptional activator in the context of low environmental oxygen. (**B**) We performed a validation experiment in which *P. aeruginosa* wild type and *anr* mutant cells were grown as microcolonies on CF airway epithelial cell monolayers. Although the validation set presents a distinct experimental system from the liquid culture based experiments in A, Node 42 activity reflected the *anr* mutant, indicating the robustness of the ADAGE model. (C) We assessed the ADAGE model on two RNA-Seq datasets both with *anr* mutant and wild type *P. aeruginosa.* Node 42 again differed (FDR q value of 0.05 in PAO1 strain and 0.10 in J215 strain) when the wild type and *anr* mutant strains were compared in both strains.

To evaluate the ADAGE model’s robustness in terms of the relationship between Anr activity and the activity of Node 42, we performed an independent experiment in which *P. aeruginosa* wild type and Δ*anr* mutant cells were grown in association with CF bronchial epithelial (CFBE) cell monolayers (Figure 4B). This experiment differed from the majority of those in the *P. aeruginosa* gene expression compendium in which the bacteria were grown as planktonic cultures. ADAGE analysis of the genome-wide expression measurements for mRNAs from both *P. aeruginosa* wild type and Δ*anr,* with three biological replicates per strain, confirmed that Node 42 reflected the absence of *anr*. This demonstrated that ADAGE not only described the patterns in data within the array experiments used to build the model, but was also able to detect these patterns in experiments performed in environments not well represented in the training dataset. ADAGE analysis also revealed that in this condition, the deletion of *anr* significantly impacted 19 other nodes (t test with FDR threshold of 0.05) consistent with the observation that Anr impacts the direct and indirect expression of many pathways in surface associated cells (41).

Since *Pseudomonas* GeneChip data were used to build the ADAGE model, and the validation experiments above employed additional microarray data, we next assessed the use of ADAGE for interpretation of RNA-Seq data. We applied the TDM (Training Distribution Matching) method described by Thompson et al. (42) to normalize RNA-Seq data to a comparable range before ADAGE analysis. Using a recently published dataset in which gene expression was analyzed in two strains and their Anr derivatives grown as colonies in an 1% oxygen atmosphere (41), we found that ADAGE’s Node 42 differed (FDR q value of 0.05 in PAO1 strain and 0.10 in J215 strain) when the wild type and Δ*anr* mutant strains were compared (Figure 4C). This demonstrates that ADAGE can also be used to interpret RNA-Seq data, and our goal is for future iterations of ADAGE to be built using both microarray and RNA-Seq data.

### ADAGE Reveals Subtle Patterns Contained in Existing Experiments

Using another previously published dataset in which *P. aeruginosa* was grown in association with CFBE cells in culture, we demonstrate that the ADAGE model can reveal important patterns associated with low magnitude gene expression changes. In this analysis, we reexamined the response of *P. aeruginosa* biofilms to challenge with either the antibiotic tobramycin or with vehicle control for thirty minutes (E-GEOD-9989, (43)). In this dataset, Nodes 39, 16, and 29 were most differentially active (Figure 5A). KEGG pathway enrichment analysis of HW genes in Nodes 16 and 29 revealed enrichment of genes involved in siderophore biosynthesis and ATPase activity, respectively (Figure 5B), and differential expression of genes in pathways upon tobramycin-treatment was evident in the array data (siderophore biosynthetic transcripts and transcripts involved in energy generation were decreased 2-29 fold and 3-20 fold respectively in response to the antibiotic treatment (see Supplemental Table 4 in Anderson et al. (43)). The third node with differential activity between tobramycin-treated and untreated cells was Node 39 (Figure 5A), and the HW genes in Node 39 were most enriched in genes associated with T3SS (Figure 5B). A standard microarray analysis approach did not detect a strong expression difference in genes involved in T3SS; among 775 differentially expressed genes (log2 fold change >1 and adjusted p value <0.05 after fitting a gene-wise linear model using the limma R package (44)), only 2 out of 18 genes in the T3SS KEGG pathway were differentially expressed (Figure 5C). However, the authors did observe a difference in T3SS-dependent cytotoxicity, and a subsequent study by the same authors demonstrated that a tobramycin-induced transcript, *mgtE,* (see Supplemental Table 3 in Anderson et al. (43)), represses *exsA,* the transcriptional activator of the T3SS (45). Thus, we propose that ADAGE can indicate meaningful differences in biology even when the transcriptional difference is low. Small differences in transcript levels associated with a differentially-active process can result from expression analysis at a time point that does not capture maximal differential expression, differential expression in only a subset of the population, or other inherent gene expression properties for transcripts of interest. Because each ADAGE node represents a multi-gene pattern that was learned from analyzing the entire expression compendium, ADAGE node-based analyses may be particularly capable of detecting subtle patterns.

**Figure 5:**
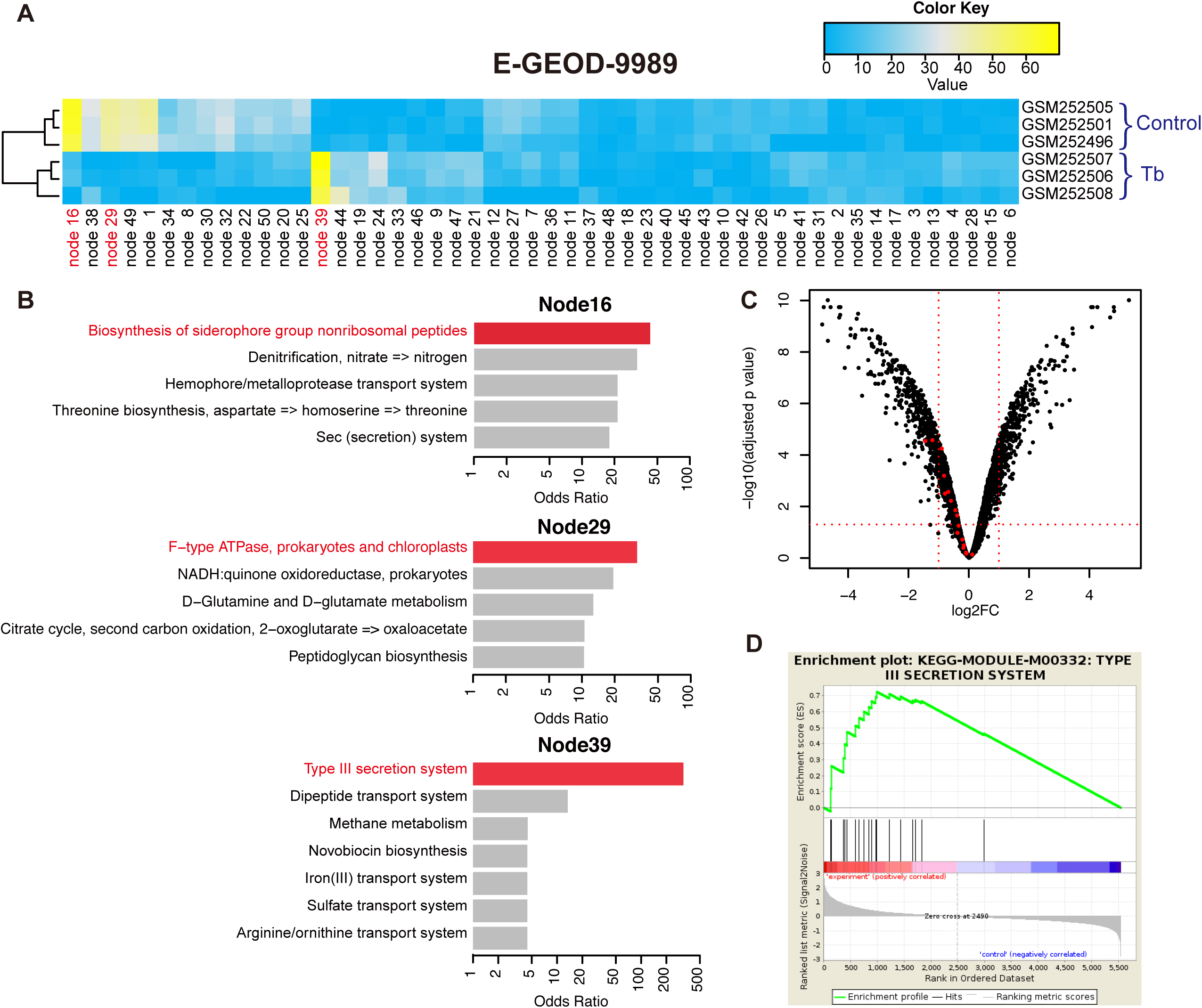
Re-analysis an existing experiment using ADAGE. (**A**) Node activity heatmap for the dataset E-GEOD-9989. Node 39, 16, 29 are the three nodes that most strongly differentiate control samples from those challenged with tobramycin. (**B**) KEGG pathway analysis of differentially active nodes. Top 5 most enriched KEGG pathways (including ties) based on odds ratios are shown for each node. Nodes 39, 16, 29 each represent a cellular process being influenced in the experiment (in red), especially Node 39, which captures the subtle change in the T3SS pathway. (C) Volcano plot showing differentially expressed genes in E-GEOD-9989. The horizontal dotted red line indicates adjusted p value cutoff of 0.05 and the two vertical lines correspond to log2 fold change of -1 and 1. Genes in the KEGG T3SS pathway (KEGG-Module-M00332) are highlighted in red. Although only two genes pass both the significance cutoff and fold change cutoff, genes in the pathway show consistently lower expression values in tobramycin treated samples. (D) Gene Set Enrichment Analysis of the T3SS pathway. GSEA also captured the consistent differential expression pattern of T3SS, which it ranked as the 12th most significantly enriched pathway.

The ability to identify meaningful small but consistent changes in gene expression is also a characteristic of GSEA analysis (46) and GSEA did identify the T3SS pathway as the 12th most significant pathway (Figure 5D). However, GSEA relies on curation to group genes together in a pathway, while ADAGE directly learns biological features from the data even in the absence of annotation. In well-studied species like *P. aeruginosa,* we were readily able to predict modulation of the T3SS pathway by looking at the identities of the HW genes of Node 39. In less well-studied species, the knowledge of genes in differentially active nodes would provide the basis for hypothesizing on the nature of involved pathways based on analysis of co-regulated genes, bioinformatics analysis of genes, and targeted genetic and biochemical studies.

### Comparison with PCA and ICA

PCA (47, 48) and ICA (49, 50) are two frequently used feature construction methods in bioinformatics that have also been applied to the analysis of gene expression data. To compare ADAGE extracted features to those from these methods, we performed analyses with each that were parallel to our ADAGE analysis. To compare with the 50 node ADAGE model, we analyzed the first 50 components in PCA and ICA applied to the expression compendium used to create ADAGE. These components explained >80% of variance in the compendium. For analysis of the DNA hybridization experiment of PAO1 and PA14 strains, we found PC4, PC5 in PCA and IC26, IC18 in ICA as the two most differentially active components in each approach (Supplementary Figure 1A). With ADAGE we analyzed only the most differentially active node, but we evaluated the top two components for PCA and ICA to avoid missing potential signals in the second most differentially active component. While many of the genes that were different in the genome hybridization comparison of strain PA14 and strain PAO1 were HW genes in PC4, PC5, IC26, and IC18, these components failed to accurately capture strain variations in other datasets (Supplementary Figure 1B). In the dataset E-GEOD-7704, *P. aeruginosa* RNA from the CF sputum was analyzed directly and RNA was harvested from the same set of sputum-derived stains after growth ex vivo. While ADAGE Node 30 differentiated the samples comprised of clinical strains from samples containing only strain PAO1, the samples containing clinical strains were heterogeneous and not uniformly different from the PAO1 samples with respect to PC4 (Figure S1) indicating that strain differences were not reliably captured by this component. Inspection of HW genes for PC4 revealed that it not only contained strain-specific genes but also a significant number of Anr-regulated genes (FDR q value = 1.0e-42), so the differential activity of PC4 is also likely to be influenced by oxygen availability; the increased activity of PC4 in the clinical strain samples from the lung samples in comparison to clinical strain samples grown in the laboratory is consistent with the finding that *P. aeruginosa* is in an oxygen limited state in the CF lung (51) (Supplementary Figure 1B). Because PCA seeks to find the direction of the largest variance, each component can become a mixture of highly variable processes that are not biologically related. While ICA decomposes data into independent signals and does not have this property, we found that the activities of strain-associated components extracted by ICA were not consistent within the replicates of individual datasets.

As in our ADAGE analyses, we also evaluated PCA and ICA components representing oxygen abundance and Anr activity. PC4 (FDR q value = 1.0e-42), PC7 (FDR q value = 4.5e-46), IC14 (FDR q value = 3.2e-20), and IC49 (FDR q value = 1.1e-19) were the components most enriched in Anr-regulated genes. While these PCA and ICA components were able to identify trends in which Anr-regulated genes were differentially expressed in response to oxygen, the resolution of the Anr signal was notably better when the ADAGE model was used when all of the experiments were considered (Supplementary Figure 2). For example, PC7 was comparable to ADAGE Node 42 in many experiments, Node 42 outperformed PC7 in the analysis of the Anr-microarray dataset, because PC7 contains other processes that also changed between wild type and *anr* mutant.

Finally, we compared the results from the analysis of E-GEOD-9989, which compared the effects of tobramycin on CFBE-associated *P.aeruginosa,* using ADAGE, PCA, and ICA. PCA agreed with ADAGE in terms of identifying changes in F-type ATPase-associated genes and transcripts associated with siderophore biosynthesis. In the ADAGE model, the node with the greatest mean difference between tobramycin-treated and untreated cultures (Node39) was most enriched in T3SS related genes (Figure 5) and this was consistent with the T3SS-dependent phenotype reported by the authors (43, 45). A similar analysis performed by PCA and ICA did not indicate changes in the T3SS pathway in the most strongly differentially active components (Supplementary Figure 3).

In summary, our comparisons with PCA and ICA showed that the biological features extracted by ADAGE were not captured clearly by either of these algorithms. While we expect that PCA and ICA captured certain other biological signals more effectively than ADAGE, our results demonstrate that ADAGE complemented these methods by identifying distinct signals.

### ADAGE Model Availability

Based on our own usage, we anticipate that our ADAGE model would provide a useful starting point for biological discovery. We have provided two example ways to leverage the model. One mode of analysis that we have demonstrated begins with the identification of differentially active nodes in a relevant dataset (e.g. the genome hybridization experiment in Figure 3 and the response to tobramycin experiment in Figure 5). By analyzing HW genes and gene pathways associated with differentially active nodes, we gained a better understanding of differences between samples and revealed the detection of subtle but consistent signals. Heatmaps showing differential node activities among samples in each experiment in the *P. aeruginosa* gene expression compendium were generated and are available at http://adage.greenelab.com/Paeruginosa-da/activity_heatmaps/index.html.

Another mode of analysis by ADAGE that we demonstrated begins with an investigator-curated list of genes related to a specific process or pathway (e.g. the Anr regulon). By examining nodes associated with the process, researchers may be able to identify novel genes associated with the process as well as other datasets in which the process is different. To facilitate such analyses, the HW genes for each node are provided in Supplementary File 3. The complete ADAGE model, for application to newly performed experiments, is available in Supplementary File 4. Software implementing ADAGE and performing all of the analyses described in this manuscript is available from https://github.com/greenelab/adage.

## Discussion

Our ADAGE method identifies biological signals (represented by nodes) by intentionally integrating noise into gene expression data prior to the data reconstruction process and model building. Thus, this method is well suited to the analysis of heterogeneous gene expression data generated in different labs, from experiments with different strains and different growth conditions. The ADAGE methodology differs from other analyses of genome-wide gene expression across large collections, which have generally been performed on homogenous collections (52, 53) or through supervised algorithms that can use known aspects of biology to separate biological signal from noise (12, 54). The *P. aeruginosa* ADAGE model, created without the use of any information on genome structure or gene functions, found that co-operonic genes and adjacent non-co-operonic genes were significantly more likely to be involved in similar processes. Furthermore, genes with similar gene-node relationships were much more likely to share KEGG function than would occur by chance. Because the building phase of ADAGE does not require any pre-specified knowledge, we anticipate that ADAGE will find use in organisms with well-curated genomes as well as in organisms for which genome curations are lacking. In organisms without curation, the ADAGE model may guide researchers towards gene sets of interest for analysis using additional computational and experimental analyses.

Analysis of existing and newly generated gene expression data using the ADAGE model found expression signatures that correlate with the comparison of different strains, and the response to low oxygen. Many genes contribute to the activity scores of each node, thus node activities can represent patterns resulting from direct or indirect aspects of a given process and may be useful in identifying patterns that are apparent in only a subset of cells in a population. Extracting these subtle but consistent changes from single experiments is difficult or impossible and requires integrative analyses leveraging information from the entire compendium.

Techniques like ADAGE, allow multi-process membership, which means that genes can be assigned to multiple distinct processes simultaneously and these processes can differ in activity independently, as is often the case in biology. In this way, ADAGE is distinct from clustering- or biclustering-based techniques such as cMonkey (55), which identify subsets of genes co-regulated in subsets of experiments. ADAGE is comparable to PCA and ICA, as each gene contributes to nodes via weight, and all nodes have a specific activity in each sample. Through comparison with PCA and ICA, we confirmed that ADAGE extracts signals distinct from these two commonly used feature construction methods. We found that PCA grouped multiple biological sources of variability into top components. Though ICA extracts independent signals, it was not able to capture the same key features of the data captured by ADAGE (strain variation and oxygen abundance). The comparison of ADAGE to PCA and ICA found that ADAGE was superior in grouping replicate samples in the same dataset, and this may reflect ADAGE’s strength in dealing with noisy measurements. We propose that both PCA and ADAGE are complementary analytical tools for the unsupervised analysis of large-scale collections of gene expression data. PCA may be preferred for a quick overview of the major sources of variations in a dataset, while ADAGE may excel in extracting differentially active biological processes.

As next generation sequencing facilitates the creation of large gene expression compendia in many organisms, algorithms capable of converting those data into insights about the underlying biological system will be required. In order to capitalize on the wealth of knowledge in large community datasets, communities need to agree upon standardized gene nomenclature for alleles across species. Alternatively, methods to extract allele information for use when two different strains are compared must be developed. In addition, the inclusion of detailed experimental information upon the deposition of gene expression data into public databases will improve the ability for community-wide data to be used by many to understand pathways and processes of interest.

We demonstrated the biological relevance of a 50-node ADAGE model, and expect that increasing node number will allow for the further separation of distinct processes with independent transcriptional signatures. Denoising autoencoders and other deep learning based methods allow for a stacked representation that maps well to layers of biological regulation. Future work will focus on building larger and deeper networks to better model complex biological systems and on incorporating multiple data types into a single model. As reviewed by leaders in the field, unsupervised use is likely to be the future of deep learning (56) and we anticipate that ADAGE and other unsupervised deep learning based approaches will continue to complement traditional feature extraction methods such as PCA and ICA in this context. We anticipate that the collection of new data, particularly data measuring new environments and genetic perturbations, will continue to refine and improve the *P. aeruginosa* ADAGE model over time.

## Materials and Methods

### Construction of a Gene Expression Compendium for P. aeruginosa

We downloaded a complete collection of *P. aeruginosa* gene expression datasets measured on the Affymetrix platform GPL84 with available supplemental CEL files from the ArrayExpress Archive of Functional Genomics Data (21) on 02/22/2014. This resulted in a collection of 109 distinct datasets covering 950 individual samples with measurements for 5549 genes. We first combined these samples generated by different laboratories into one large expression compendium using the rma function with quantile normalization provided in Bioconductor’s affy package in R (57) and the resulting expression measurements are in log_2_ scale. For autoencoder construction, we linearly transformed the expression range of each gene to be between 0 and 1. Validation datasets from the *Pseudomonas* GeneChip platform were processed concurrently through the rma function and linearly zero-one normalized using the same expression range as the compendium.

For RNA-Seq datasets, we retained genes intersecting with those existing in the compendium. The expression values of genes contained in the compendium but not measured by RNA-Seq were set to zero. To address the dynamic range differences between microarray and RNA-Seq platform, we applied the Training Distribution Matching (TDM) method to normalize RNA-Seq data and make them comparable to microarray data (42). As with the microarray validation sets, a linear zero-one normalization was performed after TDM.

### Training the ADAGE Model

We constructed a denoising autoencoder to summarize the *P. aeruginosa* gene expression compendium covering diverse genetic and environmental perturbations. We used the Theano (58) Python library to implement DA training. To train one sample, we randomly corrupted a percentage of the genes (termed the *corruption level)* by setting their input values to 0 (18). The corrupted sample x serves as input to the DA. By multiplying the corrupted sample x with a weight matrix W, we calculated the activity vector A (Formula 1). This activity vector represented the activities of each hidden node without considering the hidden bias vector b or the sigmoid transformation. To calculate the hidden representation y, we added the activity vector to b and applied a sigmoid transformation (Formula 2). Next, we computed the reconstructed input z by multiplying y with the transpose of the weight matrix W’ and adding visible bias vector b’ (Formula 3). Accurately reconstructing the input value thus represented a problem of fitting appropriate weight matrix and bias vectors to minimize the cross-entropy L_H_ between the initial input and the reconstructed input (Formula 4). To accelerate the training process, we trained the DA in batches of samples, and the number of samples in each batch was termed the *batch size*. The reconstruction error was optimized through stochastic gradient descent with the weight matrix W and bias vectors b, b’ being updated in each batch. The magnitudes of weight and bias changes were controlled by a specified *learning rate.* Training proceeded through epochs, and in each epoch training used sufficient batches to include all training samples. Training stopped once the specified number of epochs (termed the *epoch size*) was reached. A detailed description of training for denoising autoencoders has been provided by Vincent et al (18).

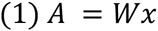

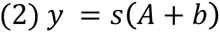

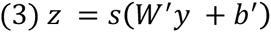

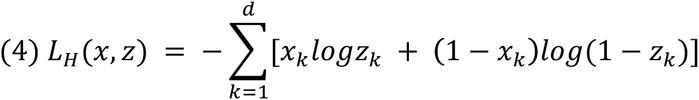

To allow the manual interpretation of nodes, we fixed the number of nodes at 50 and named them “Node##” based on the order in which they appear. We used the parameters identified as suitable for a gene expression compendium by Tan et al. (59): a batch size of 10 over 500 training epochs with a corruption level of 0.1 and a learning rate of 0.01.

After the DA was fully trained and the weight matrix was fixed, we calculated the activity value for each specific node for each specific sample in the training pool by computing the dot product of the row vector for that node in the weight matrix and the gene expression vector of the sample. We calculated the activity values of samples in newly generated validation experiments, which were not included in the training pool, in the same manner.

### Identification of High-weight Genes for each ADAGE Node

Each gene was connected to each node through a value in the weight matrix, *W*. For each node, this learned vector of weights connected that node to each gene. We calculated the standard deviation of each node’s weight vector and defined a set of high weight (HW) genes for the node that had weights two or more standard deviations away from the mean. This set of HW genes summarized the genes with the strongest influence on the node’s activity.

### Association of P. aeruginosa Operons with Specific ADAGE Nodes

To associate a specific set of operons to a node, we carried out a Gene Set Enrichment Analysis (GSEA) (46) for weight vectors. The weight vector corresponding to each constructed node was used as the weighted gene list. The curated operon information was downloaded from the Database of prOkaryotic OpeRons (DOOR) (24). We considered operons consisting of three or more genes as potential target gene sets in GSEA, and we used a false discovery rate threshold of 0.05 to identify significant associations. We calculated the overall coverage of operons as the ratio of the number of operons significantly associated with at least one node to the total number of operons curated in DOOR. Operons significantly associated with each node are provided in Supplemental File 4.

### Evaluation of the Association Between Gene Positions and ADAGE Weights

Bacterial genes are grouped by function in the genome. We tested the ADAGE model’s ability to capture such relationships in the learned weight matrix. We fitted a logistic regression model (Formula 5) with the goal of predicting whether or not a gene would be HW for a node based on two factors: the number of genes between the pair of genes (*d*) and whether or not it is co-operonic with a HW gene in the same node (*c*). We also included an interaction term between *d* and *c* in the model. We considered *d* values in the range from 1 to 10 and disregarded genes that were more than 10 genes away from the gene in question. We tested the significance of the coefficients on *d*, *c*, and the interaction term to assess the extent to which each indicated a relationship with a gene’s likelihood of being *HW* in a node.

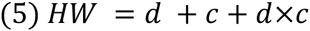

### Gene Function Assignment with the ADAGE Weight Matrix

We assessed the extent to which the ADAGE model captured genes’ functions. We employed a simple 1-nearest-neighbor classifier to assign the function of a gene. Each gene was connected to nodes through a vector of 50 weights (for a 50 node ADAGE model). For a target gene, we calculated the Euclidean distance between that gene’s weight vector and the weight vectors of all other genes. We considered the nearest neighbor to be the gene with the smallest distance. We assigned the KEGG function or functions of this closest neighbor to the target gene. To evaluate this assignment, we used KEGG pathways as the gold standards for gene function. Because one gene could be annotated with multiple KEGG pathways, we used two assessment criteria. In the first, we considered a function assignment to be correct as long as there existed an overlap between the assigned pathway and the gene’s annotated pathways. As a second evaluation, we used a more stringent definition of a correct assignment that required all of the predicted and annotated pathways to match. To evaluate the extent to which the ADAGE weight matrix captured gene functions, we compared observed accuracies with the performance of 1000 weight matrices with randomly permuted gene labels. These matrices preserve the overall weight distributions for each node but eliminate the relationship between genes and their weights. The distribution of the prediction accuracy using permuted weight matrices was plotted.

### Analysis of Sequence Divergence and Gene Expression using Affymetrix P. aeruginosa Gene Chips

To compare P *aeruginosa* wild type and Δ*anr* on airway epithelial cells, P *aeruginosa* biofilms were grown on airway epithelial cells homozygous for the *CFTR*ΔF508 mutation (CFBE41o^−^) (60) as described previously (43, 61). Data were comprised of 3 biological replicates each for wild type PAO1 and the Δ*anr* mutant. Briefly, stationary phase cultures of P *aeruginosa* grown in LB shaken at 37 °C were washed x2 and resuspended in minimal essential media (MEM) at an OD_600_=0.5. These suspensions were applied to confluent CFBE41o^−^ cells grown in 24-well plastic dishes (MatTek Corp., Ashland, MA) and incubated at 37 °C, 5% CO_2_ for 1 h, at which point planktonic cells were aspirated away and media was replaced. After an additional 90 min, planktonic cells were removed again, and the monolayer was washed x2 with PBS. Epithelial cells and attached bacterial biofilms were treated with lysozyme, and RNA was harvested using an RNeasy kit (Qiagen). RNA samples were treated with RQ1 DNAase from Promega to remove contaminating DNA, and a MICROBExpress Bacterial mRNA Enrichment Kit (Life Technologies) was used to deplete eukaryotic RNA from the samples.

For each RNA sample, cDNA samples were synthesized with Super-script III reverse transcriptase (Invitrogen, Carlsbad, CA) and NS5 primers instead of random hexamers. The cDNAs were terminally labeled with biotin-ddUTP (Enzo Bio-Array terminal labeling kit, Affymetrix) and hybridized to Affymetrix *Pseudomonas* GeneChips according to the manufacturer’s instructions with the GeneChip fluidics station 450 (Affymetrix). GeneChips were scanned with the GeneChip Scanner 3000 7G (Affymetrix) in the Dartmouth Genomics Shared Resource laboratory. The BioConductor Affy library was used to process CEL files as described above for the compendium. Data have been uploaded to GEO and are available under GSE67006.

For the genome hybridization analysis of *P. aeruginosa* strains PA14 and PAO1, genomic DNA was isolated, digested using DNAse I, denatured at 100°C for ten minutes, then labeled as described above. The GeneChips were processed as described above. Data have been uploaded to GEO and are available under GSE67038.

### Node Interpretation with GO and KEGG

We used the experimentally-derived annotations in Gene Ontology (GO) (29, 62) and KEGG pathways (25) of *P. aeruginosa* to identify the biological features captured by each node. Only terms that had more than 5 genes but fewer than 100 genes were considered. We calculated an odds ratio that indicated how over-represented each GO/KEGG term was in each node’s HW genes. Top 10 enriched pathways for some selected node are listed in Supplemental Table 1 and the full list for all 50 nodes can be downloaded from the online repository (https://github.com/greenelab/adage/blob/master/Node_interpretation/GO_KEGG_enrichment.txt).

### Sequence alignment and comparison across strains

The DNA sequences of 13 strains that have been sequenced before were obtained from the *Pseudomonas* Genome Database (63). Orthologous genes across 13 strains were aligned using Clustal Omega (64) via the EMBL-EBI webserver (65), and the alignment results including percent identity matrices and phylogenetic trees were downloaded. The phylogenetic trees in Figure 3D were drawn using Tree Graph 2 (66).

### Principal Component Analysis and Independent Component Analysis

PCA and ICA were performed in R using *prcomp* function and *fastICA* function from the fastICA package (67). For PCA we used the matrix of variable loadings as an analog to ADAGE’s weight matrix. For ICA, we used the product of the pre-whitening matrix and the estimated un-mixing matrix as the weight matrix, which first projects data onto the first 50 principal components and then projects them onto the independent components. HW genes for each component were defined in the same manner as with ADAGE: genes outside of two standard deviations of each method’s weight distribution.

### ADAGE Model and Source Code Availability

To facilitate the use of ADAGE by the *P. aeruginosa* research community, we have generated an ADAGE analysis of all of the publically-available *P. aeruginosa* gene expression experiments included in our compendium (Supplemental File 5) and provide open-source code to perform construction of ADAGE models and their application to newly generated data (https://github.com/greenelab/adage).

## Acknowledgements

This research is funded in part by the Gordon and Betty Moore Foundation’s Data-Driven Discovery Initiative through Grant GBMF4552 to CSG. JT is a Neukom Graduate Fellow supported by the William H. Neukom 1964 Institute for Computational Science. Research reported in this publication was also supported by National Institutes of Health (NIH) RO1AI091702 to DAH, T32DK007301 (Stanton) to JHH., and P30GM106394 (Stanton). This work was also supported by grants from the Cystic Fibrosis Foundation Research Development Program (STANTO07R0 to DAH and STANTO15R0 to CSG and DAH). Gene expression measurements were carried out at Genomics Shared Resource of the the Geisel School of Medicine at Dartmouth, which was established by equipment grants from the NIH and NSF and is supported in part by a Cancer Center Core Grant (P30CA023108) from the National Cancer Institute. The content is solely the responsibility of the authors and does not necessarily represent the official views of the NIH. We thank Thomas H. Hampton (Geisel School of Medicine at Dartmouth), Kevin Peterson (Dartmouth College), George O’Toole (Geisel School of Medicine at Dartmouth) and Sven D. Willger (Geisel School of Medicine at Dartmouth) for helpful discussions and assistance regarding data analysis and presentation.

## Supplemental Material

**Supplementary Figure 1:**
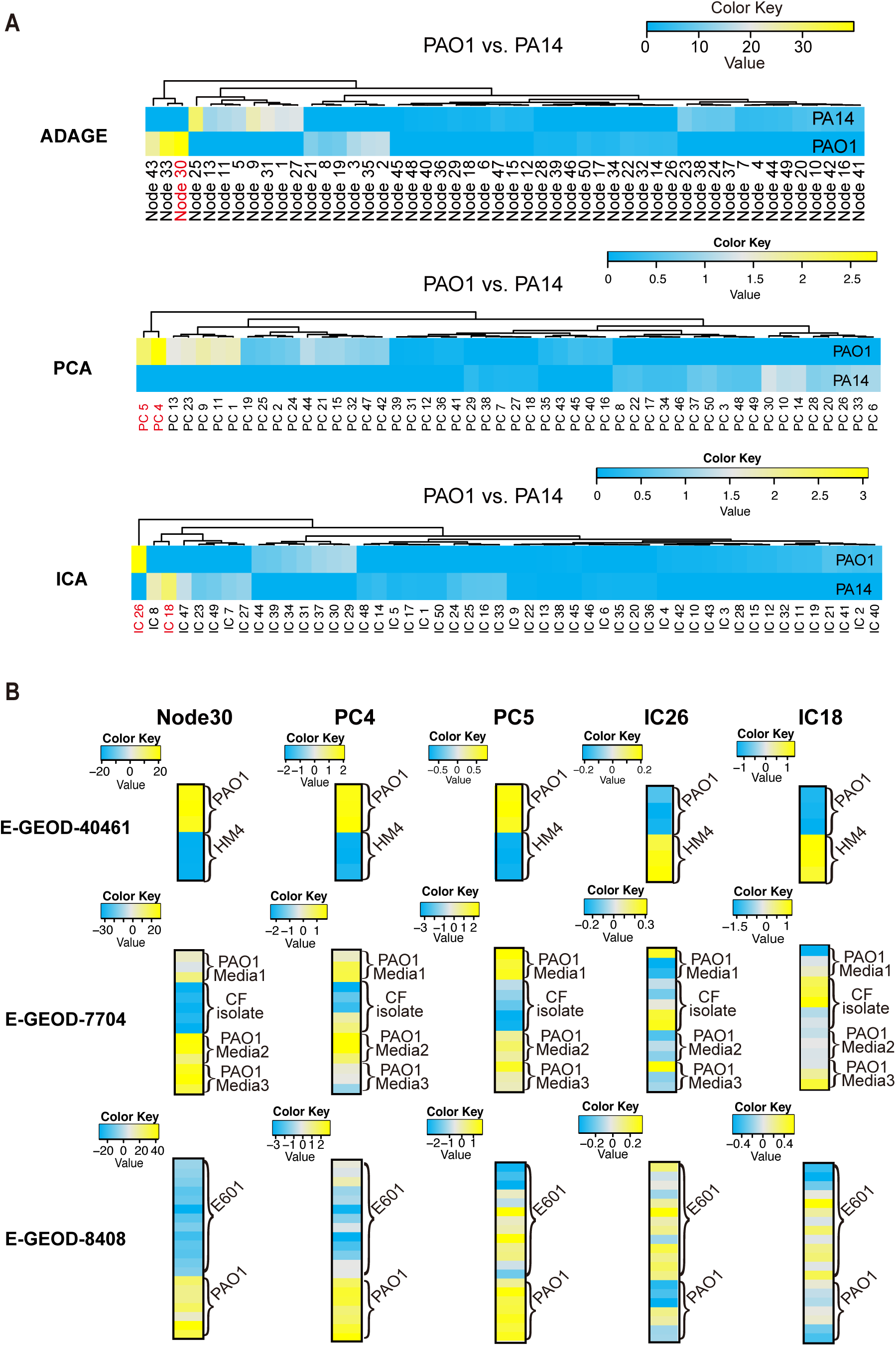
Comparison of PCA and ICA with ADAGE for the DNA hybridization experiment and extraction of strain-specific features. (**A**) In ADAGE, Node30 most differentiated PAO1 strain from PA14 strain. PC4 and PC5 in PCA and IC26 and IC18 in ICA were the components that differed the most between two strains. Top two components in PCA and ICA, as opposed to the top one component for ADAGE, were evaluated to give each method the benefit of the doubt. (**B**) Node30 from ADAGE clearly separates PAO1 strain from other strains in three independent datasets. PC4, PC5, IC26, and IC18 do not effectively capture the strain variations across the three datasets.

**Supplementary Figure 2:**
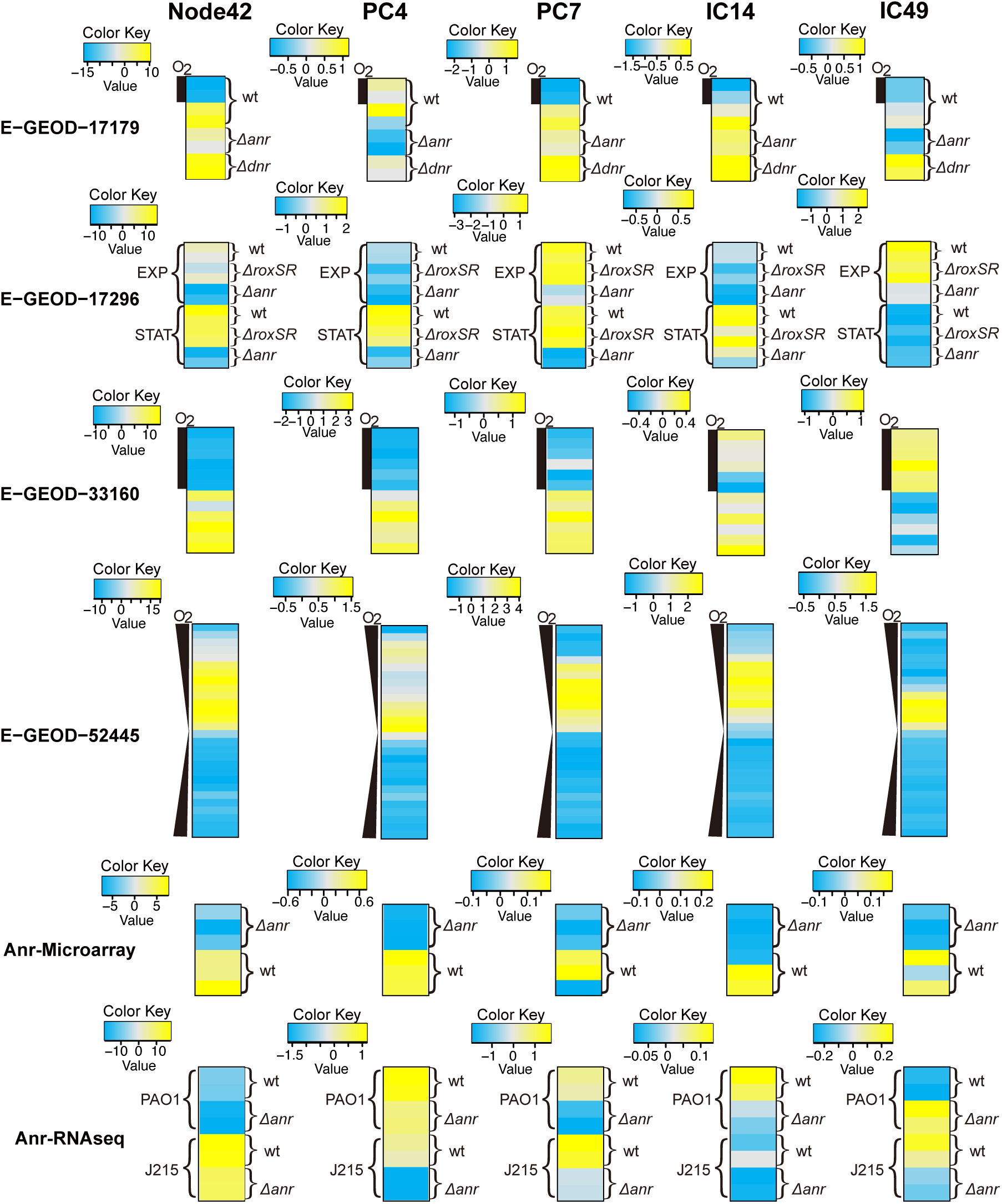
Comparison of PCA and ICA with ADAGE for the transcriptional signal of Anr and oxygen abundance. Node42 from ADAGE robustly reflects Anr activity in varying conditions including aerobic/anaerobic environment, exponential/stationary growth phase, *anr* knockouts grown on CFBE, and *anr* knockouts in PAO1 and clinical isolate (J215). PC4 does not capture Anr activity in E-GEOD-17179 and E-GEOD-17926. PC7 does not capture *anr* mutant *P. aeruginosa* grown on CFBE (the color key’s small range indicates PC7 cannot differentiate *anr* mutant from wild-type). IC14 and IC49 exhibit non-Anr patterns in multiple experiments.

**Supplementary Figure 3:**
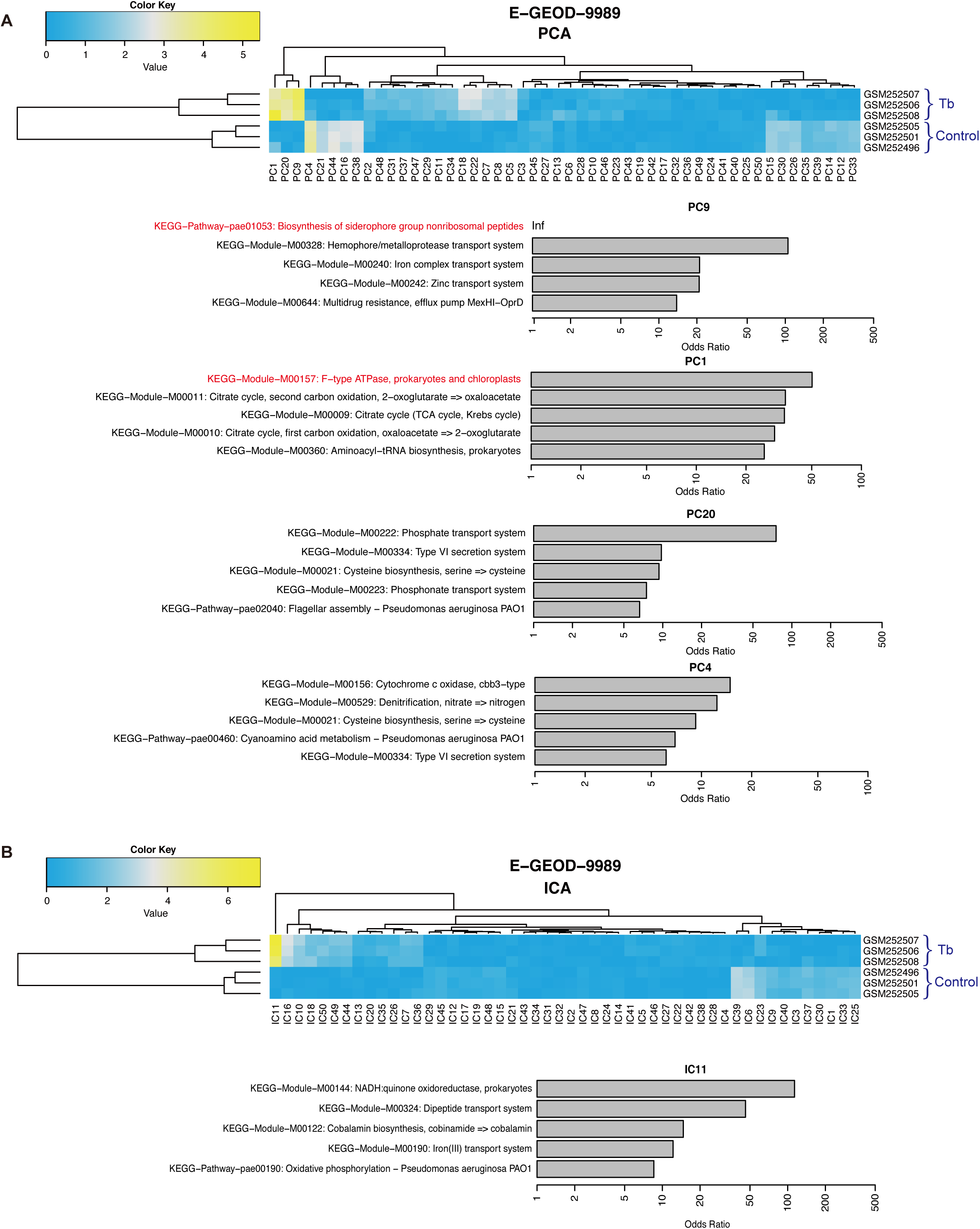
Analyzing dataset E-GEOD-9989 with PCA and ICA. (**A**) A heatmap representing principal component values shows that PC9, 1, 20, and 4 (in order of absolute difference in mean activities between two conditions) are the PCs most differentially active between samples challenged with tobramycin and controls. KEGG pathway analysis of the four PCs identified pathways known to be influenced in the dataset, such as *F-type ATPase, prokaryotes and chloroplasts* and *Biosynthesis of siderophore group nonribosomal peptides,* but did not reveal the subtle changes in the T3SS pathway. (**B**) A heatmap representing independent component values shows that IC11 is strongly differentially active in E-GEOD-9989. However, it also lacked an association with the T3SS pathway.

**Supplemental Table 1:** Top 10 associated GO terms and KEGG pathways for each node mentioned in the paper.

**Supplemental Table 2:** Anr-regulated gene list used to identify nodes significantly enriched of genes regulated by Anr.

**Supplemental File 1:** *Psuedomonas aeruginosa* gene expression compendium. This file covers all samples in publically available datasets collected before Feb. 2nd 2014. They were combined uisng the rma function with quantile normalization provided in Bioconductor’s affy R package. Only transcripts with PA IDs were maintained.

**Supplemental File 2:** ADAGE weight matrix. This file stores each gene’s weight contribution to each node.

**Supplemental File 3:** HW genes for each node and their corresponding weights in that node. Each column in this file either stores the list of high-weight genes that are outside of two standard deviations in the node’s weight distribution or stores the list of genes’ corresponding weights. High-weight gene columns are ordered by weight.

**Supplemental File 4:** Operon-node association in the ADAGE model. Significance for the association of co-operonic genes with a given node indicating that co-operonic genes are likely to have similar high weights in a node. Only significant node-operon relationships are listed. Association with a node in this analysis does not imply that the genes in the operon are necessarily HW (outside of 2 std) in that node.

**Supplemental File 5:** ADAGE node activities for each sample. The node activity is calculated as the dot product of the sample’s expression vector and the node’s weight vector.

## References

1. Schuster SC. 2008. Next-generation sequencing transforms today’s biology. Nat Methods 5:16–8.

2. Shendure J, Ji H. 2008. Next-generation DNA sequencing. Nat Biotechnol 26:1135–45.

3. van Dijk EL, Auger H, Jaszczyszyn Y, Thermes C. 2014. Ten years of next-generation sequencing technology. Trends Genet 30:418–26.

4. Vera JC, Wheat CW, Fescemyer HW, Frilander MJ, Crawford DL, Hanski I, Marden JH. 2008. Rapid transcriptome characterization for a nonmodel organism using 454 pyrosequencing. Mol Ecol 17:1636–47.

5. Surget-Groba Y, Montoya-Burgos JI. 2010. Optimization of de novo transcriptome assembly from next-generation sequencing data. Genome Res 20:1432–40.

6. Ekblom R, Galindo J. 2011. Applications of next generation sequencing in molecular ecology of non-model organisms. Heredity (Edinb) 107:1–15.

7. Cahais V, Gayral P, Tsagkogeorga G, Melo-Ferreira J, Ballenghien M, Weinert L, Chiari Y, Belkhir K, Ranwez V, Galtier N. 2012. Reference-free transcriptome assembly in non-model animals from next-generation sequencing data. Mol Ecol Resour 12:834–45.

8. Quackenbush J. 2001. Computational analysis of microarray data. Nat Rev Genet 2:418–27.

9. Brunskill EW, Aronow BJ, Georgas K, Rumballe B, Valerius MT, Aronow J, Kaimal V, Jegga AG, Yu J, Grimmond S, McMahon AP, Patterson LT, Little MH, Potter SS. 2008. Atlas of gene expression in the developing kidney at microanatomic resolution. Dev Cell 15:781–91.

10. Best JA, Blair DA, Knell J, Yang E, Mayya V, Doedens A, Dustin ML, Goldrath AW. 2013. Transcriptional insights into the CD8(+) T cell response to infection and memory T cell formation. Nat Immunol 14:404–12.

11. Tusher VG, Tibshirani R, Chu G. 2001. Significance analysis of microarrays applied to the ionizing radiation response. Proc Natl Acad Sci U S A 98:5116–21.

12. Warde-Farley D, Donaldson SL, Comes O, Zuberi K, Badrawi R, Chao P, Franz M, Grouios C, Kazi F, Lopes CT, Maitland A, Mostafavi S, Montojo J, Shao Q, Wright G, Bader GD, Morris Q. 2010. The GeneMANIA prediction server: biological network integration for gene prioritization and predicting gene function. Nucleic Acids Res 38:W214–20.

13. Greene CS, Troyanskaya OG. 2011. PILGRM: an interactive data-driven discovery platform for expert biologists. Nucleic Acids Res 39:W368–74.

14. Greene CS, Krishnan A, Wong AK, Ricciotti E, Zelaya RA, Himmelstein DS, Zhang R, Hartmann BM, Zaslavsky E, Sealfon SC, Chasman DI, FitzGerald GA, Dolinski K, Grosser T, Troyanskaya OG. 2015. Understanding multicellular function and disease with human tissue-specific networks. Nat Genet 47:569–576.

15. Tagu D, Colbourne JK, Nègre N. 2014. Genomic data integration for ecological and evolutionary traits in non-model organisms. BMC Genomics 15:490.

16. Solomon K V, Haitjema CH, Thompson DA, O’Malley MA. 2014. Extracting data from the muck: deriving biological insight from complex microbial communities and non-model organisms with next generation sequencing. Curr Opin Biotechnol 28:103–10.

17. Ranzato M, Monga R, Devin M, Chen K, Corrado G, Dean J, Le Q V., Ng AY. 2012. Building high-level features using large scale unsupervised learning, p. 81–88. *In* Proceedings of the 29th International Conference on Machine Learning (ICML-12).

18. Vincent P, Larochelle H, Bengio Y, Manzagol P-A. 2008. Extracting and composing robust features with denoising autoencoders, p. 1096–1103. *In* Proceedings of the 25th International Conference on Machine learning - ICML ’08. ACM Press, New York, New York, USA.

19. Ishii, Takaaki and Komiyama, Hiroki and Shinozaki, Takahiro and Horiuchi, Yasuo and Kuroiwa S. 2013. Reverberant Speech Recognition Based on Denoising Autoencoder, p. 3512–3516. *In* INTERSPEECH.

20. Glorot X, Bordes A, Bengio Y. 2011. Domain adaptation for large-scale sentiment classification: A deep learning approach. Proc 28th Int Conf Mach Learn.

21. Rustici G, Kolesnikov N, Brandizi M, Burdett T, Dylag M, Emam I, Farne A, Hastings E, Ison J, Keays M, Kurbatova N, Malone J, Mani R, Mupo A, Pedro Pereira R, Pilicheva E, Rung J, Sharma A, Tang YA, Ternent T, Tikhonov A, Welter D, Williams E, Brazma A, Parkinson H, Sarkans U. 2013. ArrayExpress update-trends in database growth and links to data analysis tools. Nucleic Acids Res 41:D987–90.

22. Vincent P, Larochelle H, Bengio Y, Manzagol P-A. 2008. Extracting and Composing Robust Features with Denoising AutoencodersProceedings of the 25th International Conference on Machine Learning. ACM, New York, NY, USA.

23. Vincent P, Larochelle H, Lajoie I, Bengio Y, Manzagol P-A. 2010. Stacked denoising autoencoders: Learning useful representations in a deep network with a local denoising criterion. J Mach Learn Res 11:3371–3408.

24. Mao X, Ma Q, Zhou C, Chen X, Zhang H, Yang J, Mao F, Lai W, Xu Y. 2014. DOOR 2.0: presenting operons and their functions through dynamic and integrated views. Nucleic Acids Res 42:D654–9.

25. Kanehisa M. 2000. KEGG: Kyoto Encyclopedia of Genes and Genomes. Nucleic Acids Res 28:27–30.

26. Lee DG, Urbach JM, Wu G, Liberati NT, Feinbaum RL, Miyata S, Diggins LT, He J, Saucier M, Déziel E, Friedman L, Li L, Grills G, Montgomery K, Kucherlapati R, Rahme LG, Ausubel FM. 2006. Genomic analysis reveals that *Pseudomonas aeruginosa* virulence is combinatorial. Genome Biol 7:R90.

27. Watson JM, Holloway BW. 1976. Suppressor mutations in *Pseudomonas aeruginosa*. J Bacteriol 125:780–6.

28. Rahme LG, Stevens EJ, Wolfort SF, Shao J, Tompkins RG, Ausubel FM. 1995. Common virulence factors for bacterial pathogenicity in plants and animals. Science 268:1899–902.

29. Harris MA, Clark J, Ireland A, Lomax J, Ashburner M, Foulger R, Eilbeck K, Lewis S, Marshall B, Mungall C, Richter J, Rubin GM, Blake JA, Bult C, Dolan M, Drabkin H, Eppig JT, Hill DP, Ni L, Ringwald M, Balakrishnan R, Cherry JM, Christie KR, Costanzo MC, Dwight SS, Engel S, Fisk DG, Hirschman JE, Hong EL, Nash RS, Sethuraman A, Theesfeld CL, Botstein D, Dolinski K, Feierbach B, Berardini T, Mundodi S, Rhee SY, Apweiler R, Barrell D, Camon E, Dimmer E, Lee V, Chisholm R, Gaudet P, Kibbe W, Kishore R, Schwarz EM, Sternberg P, Gwinn M, Hannick L, Wortman J, Berriman M, Wood V, de la Cruz N, Tonellato P, Jaiswal P, Seigfried T, White R. 2004. The Gene Ontology (GO) database and informatics resource. Nucleic Acids Res 32:D258–61.

30. Mooij MJ, Drenkard E, Llamas MA, Vandenbroucke-Grauls CMJE, Savelkoul PHM, Ausubel FM, Bitter W. 2007. Characterization of the integrated filamentous phage Pf5 and its involvement in small-colony formation. Microbiology 153:1790–8.

31. Köhler T, Donner V, van Delden C. 2010. Lipopolysaccharide as shield and receptor for R-pyocin-mediated killing in *Pseudomonas aeruginosa*. J Bacteriol 192:1921–8.

32. Son MS, Matthews WJ, Kang Y, Nguyen DT, Hoang TT. 2007. In vivo evidence of *Pseudomonas aeruginosa* nutrient acquisition and pathogenesis in the lungs of cystic fibrosis patients. Infect Immun 75:5313–24.

33. Tralau T, Vuilleumier S, Thibault C, Campbell BJ, Hart CA, Kertesz MA. 2007. Transcriptomic analysis of the sulfate starvation response of *Pseudomonas aeruginosa*. J Bacteriol 189:6743–50.

34. Kang Y, Nguyen DT, Son MS, Hoang TT. 2008. The *Pseudomonas aeruginosa* PsrA responds to long-chain fatty acid signals to regulate the *fadBA5* beta-oxidation operon. Microbiology 154:1584–98.

35. Zimmermann A, Reimmann C, Galimand M, Haas D. 1991. Anaerobic growth and cyanide synthesis of *Pseudomonas aeruginosa* depend on *anr*, a regulatory gene homologous with *fnr* of *Escherichia coli*. Mol Microbiol 5:1483–90.

36. Jackson AA, Gross MJ, Daniels EF, Hampton TH, Hammond JH, Vallet-Gely I, Dove SL, Stanton BA, Hogan DA. 2013. Anr and its activation by PlcH activity in *Pseudomonas aeruginosa* host colonization and virulence. J Bacteriol 195:3093–104.

37. Trunk K, Benkert B, Quäck N, Münch R, Scheer M, Garbe J, Jänsch L, Trost M, Wehland J, Buer J, Jahn M, Schobert M, Jahn D. 2010. Anaerobic adaptation in *Pseudomonas aeruginosa*: definition of the Anr and Dnr regulons. Environ Microbiol 12:1719–33.

38. Kawakami T, Kuroki M, Ishii M, Igarashi Y, Arai H. 2010. Differential expression of multiple terminal oxidases for aerobic respiration in *Pseudomonas aeruginosa*. Environ Microbiol 12:1399–412.

39. Tielen P, Rosin N, Meyer A-K, Dohnt K, Haddad I, Jänsch L, Klein J, Narten M, Pommerenke C, Scheer M, Schobert M, Schomburg D, Thielen B, Jahn D. 2013. Regulatory and metabolic networks for the adaptation of *Pseudomonas aeruginosa* biofilms to urinary tract-like conditions. PLoS One 8:e71845.

40. He FQ, Wang W, Zheng P, Sudhakar P, Sun J, Zeng A-P. 2014. Essential O2-responsive genes of *Pseudomonas aeruginosa* and their network revealed by integrating dynamic data from inverted conditions. Integr Biol (Camb) 6:215–23.

41. Hammond JH, Dolben EF, Smith TJ, Bhuju S, Hogan DA. 2015. Links between Anr and quorum sensing in *Pseudomonas aeruginosa* biofilms. J Bacteriol 197:2810–20.

42. Thompson JA, Tan J, Greene CS. 2015. Cross-platform normalization of microarray and RNA-seq data for machine learning applications.

43. Anderson GG, Moreau-Marquis S, Stanton BA, O’Toole GA. 2008. In vitro analysis of tobramycin-treated *Pseudomonas aeruginosa* biofilms on cystic fibrosis-derived airway epithelial cells. Infect Immun 76:1423–33.

44. Ritchie ME, Phipson B, Wu D, Hu Y, Law CW, Shi W, Smyth GK. 2015. limma powers differential expression analyses for RNA-sequencing and microarray studies. Nucleic Acids Res gkv007-.

45. Anderson GG, Yahr TL, Lovewell RR, O’Toole GA. 2010. The *Pseudomonas aeruginosa* magnesium transporter MgtE inhibits transcription of the type III secretion system. Infect Immun 78:1239–49.

46. Subramanian A, Tamayo P, Mootha VK, Mukherjee S, Ebert BL, Gillette MA, Paulovich A, Pomeroy SL, Golub TR, Lander ES, Mesirov JP. 2005. Gene set enrichment analysis: a knowledge-based approach for interpreting genome-wide expression profiles. Proc Natl Acad Sci U S A 102:15545–50.

47. Misra J, Schmitt W, Hwang D, Hsiao L-L, Gullans S, Stephanopoulos G, Stephanopoulos G. 2002. Interactive exploration of microarray gene expression patterns in a reduced dimensional space. Genome Res 12:1112–20.

48. Fehrmann RSN, Karjalainen JM, Krajewska M, Westra H-J, Maloney D, Simeonov A, Pers TH, Hirschhorn JN, Jansen RC, Schultes EA, van Haagen HHHBM, de Vries EGE, Te Meerman GJ, Wijmenga C, van Vugt MATM, Franke L. 2015. Gene expression analysis identifies global gene dosage sensitivity in cancer. Nat Genet 47:115–125.

49. Lee S-I, Batzoglou S. 2003. Application of independent component analysis to microarrays. Genome Biol 4:R76.

50. Engreitz JM, Daigle BJ, Marshall JJ, Altman RB. 2010. Independent component analysis: mining microarray data for fundamental human gene expression modules. J Biomed Inform 43:932–44.

51. Worlitzsch D, Tarran R, Ulrich M, Schwab U, Cekici A, Meyer KC, Birrer P, Bellon G, Berger J, Weiss T, Botzenhart K, Yankaskas JR, Randell S, Boucher RC, Döring G. 2002. Effects of reduced mucus oxygen concentration in airway *Pseudomonas* infections of cystic fibrosis patients. J Clin Invest 109:317–25.

52. The Cancer Genome Atlas Network. 2012. Comprehensive molecular portraits of human breast tumours. Nature 490:61–70.

53. The Cancer Genome Atlas Network. 2011. Integrated genomic analyses of ovarian carcinoma. Nature 474:609–615.

54. Park CY, Wong AK, Greene CS, Rowland J, Guan Y, Bongo LA, Burdine RD, Troyanskaya OG. 2013. Functional knowledge transfer for high-accuracy prediction of under-studied biological processes. PLoS Comput Biol 9:e1002957.

55. Reiss DJ, Baliga NS, Bonneau R. 2006. Integrated biclustering of heterogeneous genome-wide datasets for the inference of global regulatory networks. BMC Bioinformatics 7:280.

56. LeCun Y, Bengio Y, Hinton G. 2015. Deep learning. Nature 521:436–444.

57. Gautier L, Cope L, Bolstad BM, Irizarry RA. 2004. affy-analysis of Affymetrix GeneChip data at the probe level. Bioinformatics 20:307–15.

58. J. Bergstra, O. Breuleux, F. Bastien, P. Lamblin, R. Pascanu, G. Desjardins, J. Turian DW-F and YB. 2010. Theano: A CPU and GPU Math Expression CompilerProceedings of the Python for Scientific Computing Conference (SciPy) 2010. June 30 - July 3, Austin, TX.

59. Tan J, Ung M, Cheng C, Greene CS. 2015. Unsupervised feature construction and knowledge extraction from genome-wide assays of breast cancer with denoising autoencoders. Pac Symp Biocomput 20:132–43.

60. Cozens AL, Yezzi MJ, Chin L, Simon EM, Finkbeiner WE, Wagner JA, Gruenert DC. 1992. Characterization of immortal cystic fibrosis tracheobronchial gland epithelial cells. Proc Natl Acad Sci U S A 89:5171–5.

61. Moreau-Marquis S, Bomberger JM, Anderson GG, Swiatecka-Urban A, Ye S, O’Toole GA, Stanton BA. 2008. The DeltaF508-CFTR mutation results in increased biofilm formation by *Pseudomonas aeruginosa* by increasing iron availability. Am J Physiol Lung Cell Mol Physiol 295:L25–37.

62. Ashburner M, Ball CA, Blake JA, Botstein D, Butler H, Cherry JM, Davis AP, Dolinski K, Dwight SS, Eppig JT, Harris MA, Hill DP, Issel-Tarver L, Kasarskis A, Lewis S, Matese JC, Richardson JE, Ringwald M, Rubin GM, Sherlock G. 2000. Gene ontology: tool for the unification of biology. The Gene Ontology Consortium. Nat Genet 25:25–9.

63. Winsor GL, Lam DKW, Fleming L, Lo R, Whiteside MD, Yu NY, Hancock REW, Brinkman FSL. 2011. Pseudomonas Genome Database: improved comparative analysis and population genomics capability for *Pseudomonas* genomes. Nucleic Acids Res 39D596–600:.

64. Sievers F, Wilm A, Dineen D, Gibson TJ, Karplus K, Li W, Lopez R, McWilliam H, Remmert M, Söding J, Thompson JD, Higgins DG. 2011. Fast, scalable generation of high-quality protein multiple sequence alignments using Clustal Omega. Mol Syst Biol 7:539.

65. McWilliam H, Li W, Uludag M, Squizzato S, Park YM, Buso N, Cowley AP, Lopez R. 2013. Analysis Tool Web Services from the EMBL-EBI. Nucleic Acids Res 41:W597–600.

66. Stöver BC, Müller KF. 2010. TreeGraph 2: combining and visualizing evidence from different phylogenetic analyses. BMC Bioinformatics 11:7.

67. Marchini JL, Heaton C, Ripley BD. 2013. fastICA: FastICA Algorithms to perform ICA and Projection Pursuit.

